# Improving sequence alignments with AlphaFold2 regardless of structural modeling accuracy

**DOI:** 10.1101/2022.05.24.492699

**Authors:** Athanasios Baltzis, Leila Mansouri, Suzanne Jin, Björn E. Langer, Ionas Erb, Cedric Notredame

## Abstract

Protein sequence alignments are essential to structural, evolutionary and functional analysis but their accuracy is often limited by sequence similarity unless molecular structures are available. Protein structures predicted at experimental grade accuracy, as achieved by AlphaFold2, could therefore have a major impact on sequence analysis. Here, we find that multiple sequence alignments estimated on AlphaFold2 predictions are almost as accurate as alignments estimated on experimental structures and significantly superior to sequence-based alignments. We also show that AlphaFold2 structural models of relatively low quality can be used to obtain highly accurate alignments. These results suggest that, besides structure modeling, AlphaFold2 encodes higher-order dependencies that can be exploited for sequence analysis.

## Main Text

Protein multiple sequence alignment (MSA) is the most widely used modeling technique in biology^1^. Its many applications include structural, functional, and evolutionary analyses^2,3^. Their computation typically relies on amino-acid substitution matrices and only achieves sufficient levels of accuracy when comparing sequences that are more than 20% identical^4^. Alignments based on structural comparisons are, by contrast, much less sensitive to low sequence identity levels and have routinely been employed as standards of truth when evaluating sequence alignment algorithms^5^. Thus, having more high quality structural data associated with sequences would substantially improve protein sequence alignments, and therefore expand their usability.

While the scarcity of experimental structural data currently limits the use of structure-based sequence alignments, recent results indicate that inferring protein structure using deep learning techniques is becoming increasingly effective, with claims that AlphaFold2 (AF2)^6^ can predict structures at angstrom-level accuracy for most proteins^7,8,9^. Furthermore, other work indicates that predicted protein folding code elements can be used to improve sequence homology detection^10^. Given these advances, we asked whether AF2 models can be used to compute highly accurate protein MSAs.

To validate the potential of AF2 models for the generation of accurate structure-based sequence alignments, we filtered the PDB^11^ to select protein sequences that were not part of the AF2 training set (see Methods). This highly stringent process left us with a total of 153 proteins that can be associated with 12 PFAM^12^ families. Sequences within each family were multiply aligned using: (1) experimental structures (MSA-PDB); (2) predicted structures (MSA-AF2); or (3) sequence information only (MSA-Seq). We used MSA-PDB as a reference to assess the alignment accuracy of every pair of sequences within MSA-Seq and MSA-AF2. Accuracy was estimated using the Sum of Pairs (SoP) measure, which measures the fraction of pairs of residues identically aligned across two alternative alignments of the same sequences. Our results indicate that MSA-AF2 is clearly superior to sequence-based alignments, with 78% of the sequence pairs being more accurately aligned in MSA-AF2 than in MSA-Seq (Fig. 1a). Across all MSAs, the difference is highly significant, with an average improvement by 23.62 percent points of MSA-AF2 over MSA-Seq (93.95% vs 70.33%, Table 1, Supplementary Table 1). This difference is particularly clear in datasets containing repeated elements, a feature known to compromise most sequence alignment procedures (e.g. PF13306, Supplementary Fig. 1). The high levels of accuracy measured on MSA-AF2 alignments confirm that the quality of these alignments is comparable to their reference^13^. These results support the notion that AF2 predicted structures could be systematically used to replace sequence information.

**Table 1.**
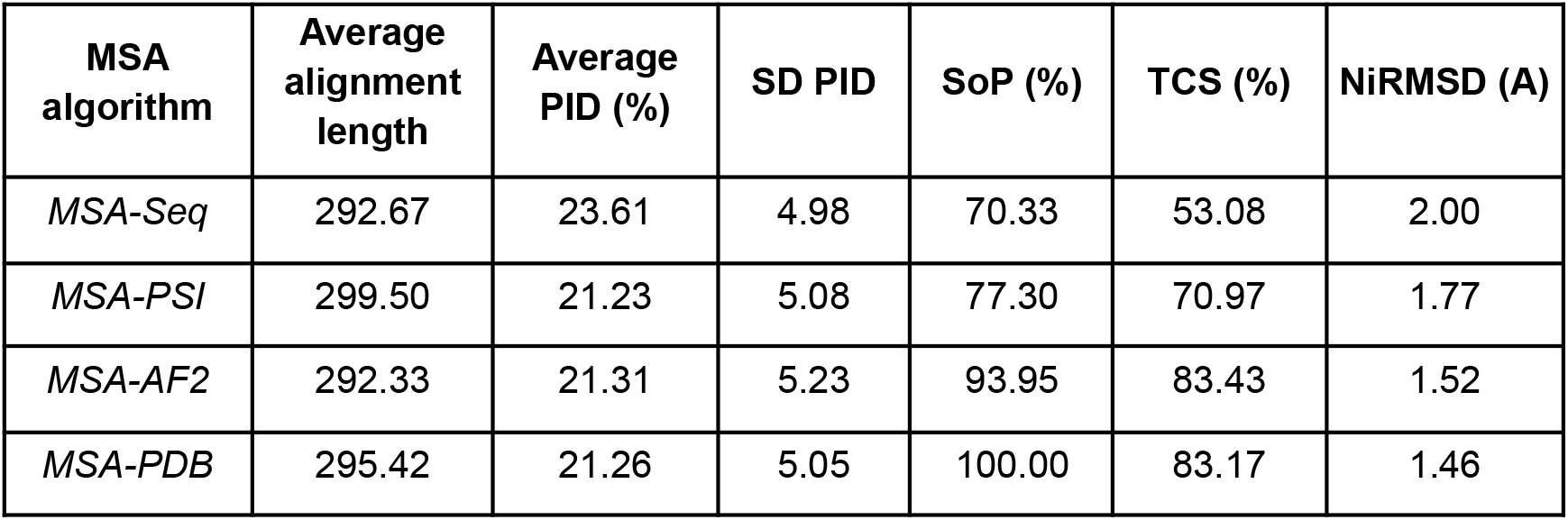
Average alignment length, Percent Identity (PID) with its standard deviation, SoP, TCS and NiRMSD scores per MSA algorithm over the 12 datasets.

| **MSA algorithm** | **Average alignment length** | **Average PID (%)** | **SD PID** | **SoP (%)** | **TCS (%)** | **NiRMSD (A)** |
| --- | --- | --- | --- | --- | --- | --- |
| *MSA-Seq* | 292.67 | 23.61 | 4.98 | 70.33 | 53.08 | 2.00 |
| *MSA-PSI* | 299.50 | 21.23 | 5.08 | 77.30 | 70.97 | 1.77 |
| *MSA-AF2* | 292.33 | 21.31 | 5.23 | 93.95 | 83.43 | 1.52 |
| *MSA-PDB* | 295.42 | 21.26 | 5.05 | 100.00 | 83.17 | 1.46 |

**Fig. 1.**
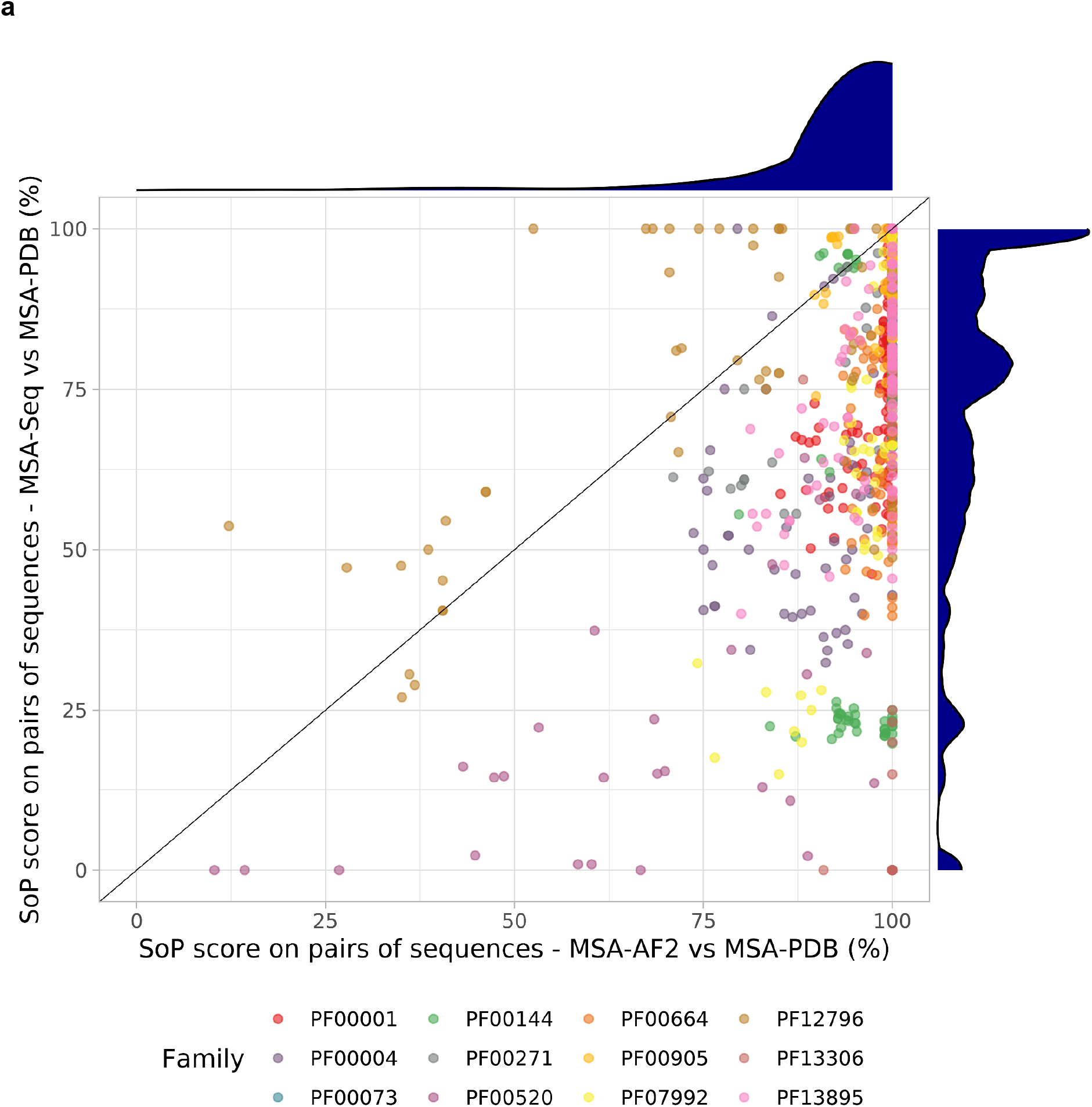
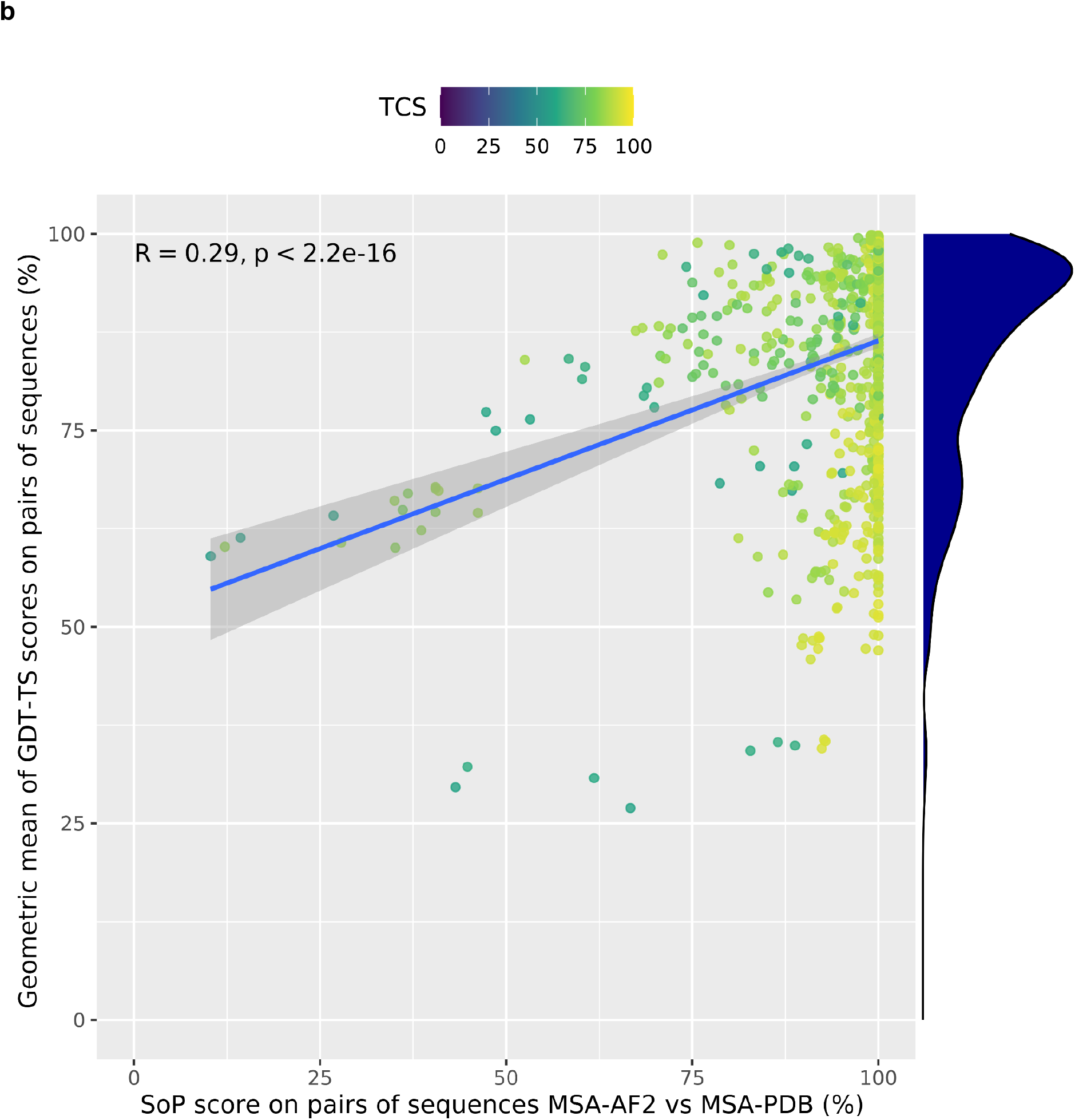
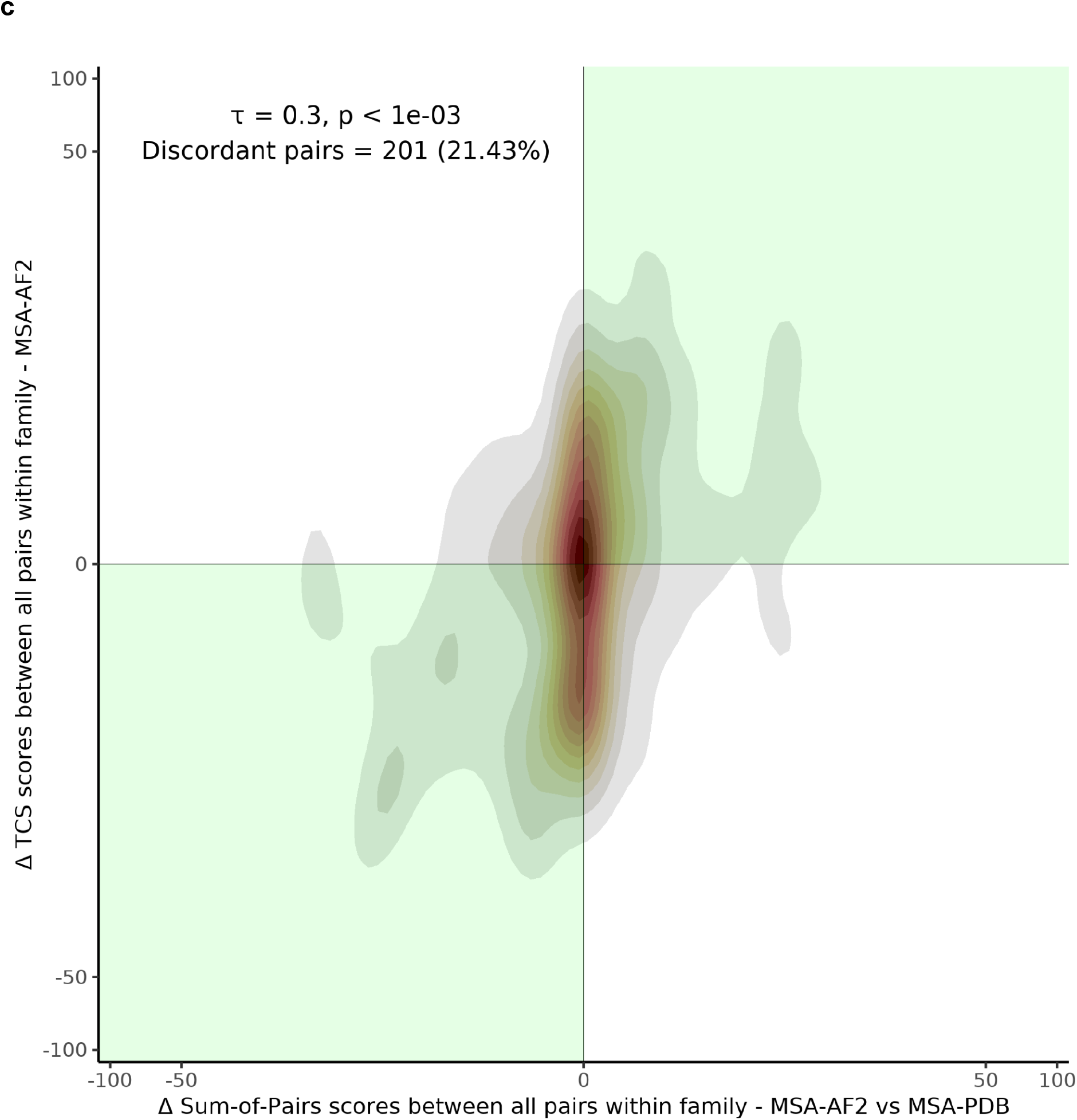
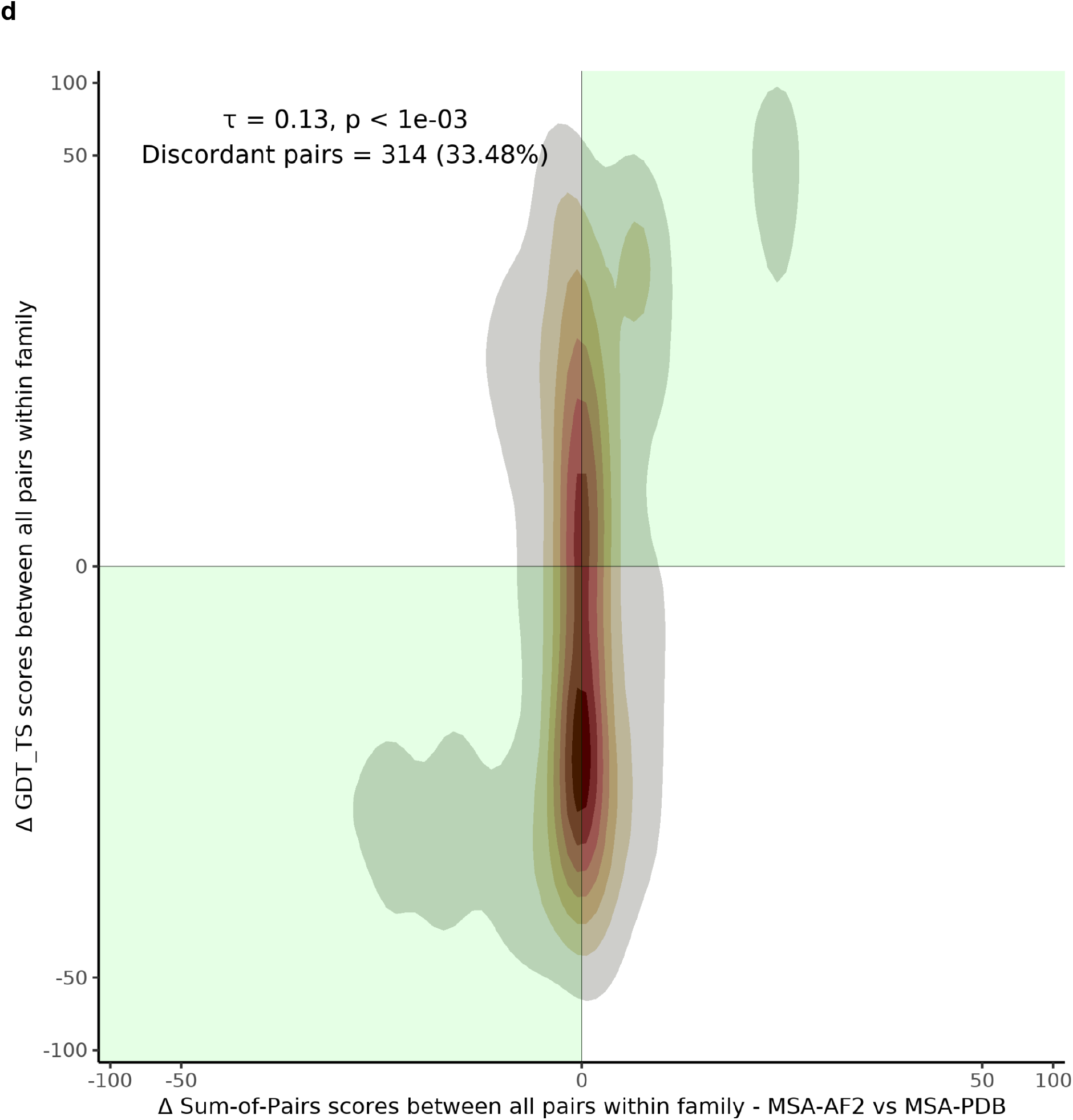
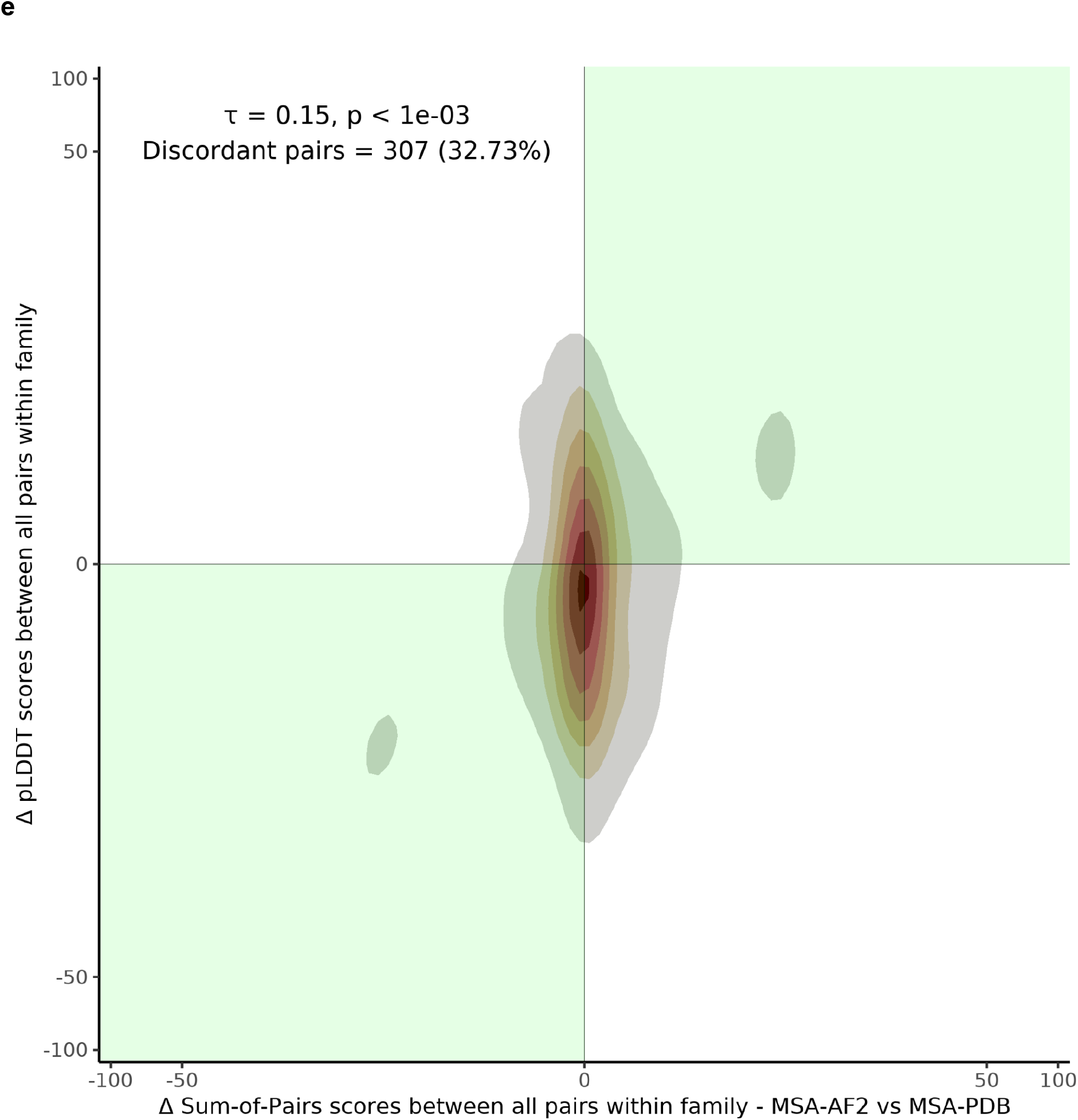
Performance of structure-based multiple sequence alignments using AlphaFold2 models (MSA-AF2). **a**, Comparison of alignment accuracy of MSAs based on sequence information (MSA-Seq, y-axis) and on predicted structural information (MSA-AF2, x-axis). Accuracy is measured against the reference MSAs based on experimental structural data (MSA-PDB) using the Sums-of-Pairs (SoP) measure estimated of every pair of sequences within the respective MSA. Points are colored by dataset and the marginal density plots represent their distribution across the considered axis. **b**, Comparison between the geometric mean of the model structural correctness (GDT_TS) and alignment accuracy (SoP) for every pair of aligned sequences within the MSA-AF2 MSAs. Points are colored by the geometric mean of the predicted sequence accuracy (TCS) for each pair. The marginal density plot represents the distribution of the number of pairs along the considered axis. The blue line represents the Pearson correlation (R=0.29 and p-value=2.2e-16) as provided by the ggpubr package. **c-e**, log/log representation of discordance between predicted sequence alignment accuracy (TCS) (c), model structural accuracy (GDT-TS) (d) or predicted model structural accuracy (pLDDT) (e) and the equivalent differences in sequence accuracy (SoP) scores when comparing MSA-AF2 against MSA-PDB. The green shaded areas correspond to the 1st and 3rd quadrants. The relationship between the two variables is indicated by a Kendall correlation coefficient (τ) and its assigned p-value. The text label also includes the number and percentage of discordant pairs over all the given pairs.

To determine whether AF2 performance is the main driver of sequence alignment accuracy (Supplementary Fig. 2), we compared sequence alignment and structural prediction accuracies. We did so by measuring the alignment score of every pair of sequences as they appear in their respective MSAs (SoP) versus the average correctness of the corresponding AF2 structures (GDT-TS geometric mean). The results paint a more complex picture than anticipated. Many aligned pairs achieve high alignment accuracy despite being based on AF2 models scoring lower than the 75% GDT-TS threshold required for atomic resolution precision^14^. Specifically, 80.7% of such pairs give rise to AF2-based alignments that are superior to their sequence-based counterparts. Altogether, these pairs of lower correctness structures occur in 18.3% of all the 938 possible pairs of sequences (Supplementary Table 2). Given that the structure-based alignments of sequence pairs rely only on C-alpha superpositions and ignore sequence information (see Methods), it is unclear why so many incorrect AF2 models contribute to highly accurate sequence alignments.

One possible explanation is that some of the AF2 predictions, albeit different from the experimental reference (i.e. low GDT-TS), may preserve amino-acid homologous relationships across protein family members. This preservation could make the AF2 structures easier to superpose and would therefore lead to more consistent alignments (see Methods). We quantified this effect using the Transitive Consistency Score (TCS)^15^. Our results indicate that MSA-AF2 alignments featuring low GDT-TS sequences can display a wide range of TCS scores (gradient overlay Fig. 1b). Overall the 22.8% pairs with a GDT-TS lower than 75% have TCS scores ranging between 54.5% and 95.5%. The very existence of low GDT-TS/high TCS alignments confirms that some of the incorrectly predicted structures are sufficiently superposable to yield consistent alignments. Their relatively high accuracy shows the benefits of such predictions for sequence analysis.

These results appear to be at odds with the expectation that structural correctness measures such as GDT-TS and the AF2 prediction reliability index, pLDDT, should be the best predictors of accuracy for alignments evaluated against structure-based references. To analyze this effect further, we compared the capacity of GDT-TS, pLDDT and TCS to discriminate between high and low accuracy alignments. We did so by quantifying the average SoP of each sequence within its MSA and by estimating its concordance with each of the three measures. When running a Kendall correlation analysis within families only (see Methods), we observed considerably less discordance between TCS and SoP (21.43% of discordant pairs, Kendall τ = 0.3, Fig. 1c) than for GDT-TS (33.48%, τ = 0.13, Fig. 1d) or pLDDT (32.73%, τ = 0.15, Fig. 1e) (Supplementary Table 3). This indicates that when aligning AF2 structures, consistency, as estimated by the TCS, is a much better predictor of alignment accuracy than the estimators of structural correctness.

The high accuracy of the MSA-AF2 alignments, regardless of the correctness of their predicted structures, may also indicate the AF2 algorithm’s capacity to integrate structural and evolutionary data. Indeed, the training of AF2 involves combining the information content of experimentally determined structures along with MSAs consisting of large compilations of naturally occurring mutations selected for maintaining a functional fold. Evolutionary information is well established as a means to improve sequence alignments^16,17^. For instance, PSI-Coffee encompasses evolutionary information by replacing every sequence with a position specific scoring scheme, often referred to as a profile. This provides a powerful way to average information content across protein family members but it discards any information associated with covariation between sites. When we applied the PSI-Coffee profile-based approach to our dataset, it systematically produced MSAs of intermediate accuracy between sequence and AF2-based alignments (Table 1, Supplementary Table 1). This observation suggests that the way AF2 models incorporate evolutionary information possibly accounts for some aspects of covariation between sites.

We showed here that the structural models generated using AF2 lead to alignments superior to the ones achieved using the best profile-based solution^18^. We hypothesize that this improvement results from deep networks identifying higher-order relationships among sites thus allowing for more accurate alignments than simpler methods like PSI-Coffee. The robustness of AF2 sequence alignments regardless of the accuracy of the predicted structures opens the way for their systematic use in any procedure requiring accurate sequence alignments.

## Methods

### Reference datasets

We assembled a structure-based dataset suitable for AF2^6^ predictions, highly discriminative and independent of any AF2 training set. This was achieved by collecting all the 23,300 Protein Data Bank (PDB)^11^ entries (07/2020 release) with a release date posterior to AF2 (04/2018). These entries were assigned to PFAM^12^ families (release 28) using HMMER3/f (Version 3.1b1 | May 2013) with default parameters (E-value threshold set at 10.0). This procedure gave rise to 39,476 domain segments longer than 80 AA that were turned into a non-redundant dataset featuring 3,419 segments (CD-HIT^19,20^ version 4.8.1, threshold 70% sequence identity), each unambiguously assigned to a PFAM family. We grouped all the segments labeled with the same PFAM family and kept the 31 datasets featuring 10 or more segments (461 segments in total). In order to generate a final test set as compact and informative as possible, we removed the 16 datasets for which the agreement between the sequence MSA and the reference structure-based MSA was higher than 75%. We also removed two datasets featuring sequences longer than 400 amino acids and one dataset containing a multi-domain protein. On top of this, we discarded two segments from the PF00520 dataset whose very short length suggests truncation. Overall this procedure led to a test set featuring 153 PDB chains belonging to 12 distinct PFAM families and comprising between 8 and 16 sequences each (Supplementary Table 1). The complete list of all the selected segments, including the discarded ones, is available from Zenodo (https://doi.org/10.5281/zenodo.6564253).

### Sequence and Structure-based Multiple Sequence Alignments

On the basis of recent benchmarks^13,21^, five different sequence-based MSA algorithms (FAMSA v.1.6.218^21^, MAFFT G-INS-i v.7.45319^22^, MSAProbs v.0.9.720^23^, T-Coffee v.13.45.57.844f401^24^, PSI-Coffee v.13.45.57.844f401^16^) and three structure-based MSA algorithms (3D-Coffee_SAP+TMalign v.13.45.57.844f401^25^, 3D-Coffee_TMalign v.13.45.57.844f401^25^, mTM-align v.20180725^26^) were tested.

The sequences of the PDB-derived structures were multiply aligned using either structure-based or sequence-based alignment algorithms and the best algorithms for sequences or structures respectively were selected on the basis of their average structural correctness using Normalized intra-molecular Root Mean Squared Deviation (NiRMSD, Supplementary Table 4), a normalized distance-based-RMSD^27^ that compares intramolecular distance variations to assess the structural correctness of sequence alignments. 3D-Coffee_SAP+TMalign, the best performing structure-based MSA algorithm, was used as a reference method to perform all the structural MSAs using either PDB or AF2 structures (MSA-PDB and MSA-AF2 respectively). The best sequence-based MSA alignment algorithm was MAFFT G-INS-i which was used for MSA-Seq. The PSI-Coffee alignments (MSA-PSI) were assembled using the psicoffee mode of T-Coffee (v.13.45.57.844f401,`-mode=psicoffee`) where we fetched for each input sequence the 100 most informative homologs by conducting a BLAST search against UniRef50. The collected BLAST alignments were stacked onto the corresponding query sequence, such that the result could be used as profile templates when generating the T-Coffee pairwise libraries^28^ used to produce a regular T-Coffee MSA.

### De novo structure prediction and evaluation

Structure predictions were carried out using AlphaFold2 (AF2) v.2.0.0^6^ with the default parameters:

~~~
-full_dbs preset -max_template_date 2020-05-14
~~~

AF2 provides for each predicted structure a reliability index named pLDDT that reports the fraction of residues expected to be correctly predicted. Furthermore, the correctness of the predicted structures was estimated via the Global Distance Test Total Score (GDT-TS, TM-Score^29^ package), one of the standard CASP measures. Given an experimental structure (PDB-derived) and its corresponding prediction, GDT-TS reports the fraction of C_β_ (C_α_ in case of glycines) superimposed below different distance thresholds after a rigid superposition.

### Alignment accuracy measures

The sequence-based alignments (MSA-Seq, MSA-PSI) and the structure-based alignments estimated on AF2 predicted models (MSA-AF2) were compared with the structure-based MSAs computed on experimental structures (MSA-PDB) and evaluated using the Sums of Pairs (SoP) measure as implemented in the T-Coffee package (Version_13.45.57.844f401). The SoP reports the fraction of aligned pairs occurring in the reference MSA that are recovered in the evaluated one. The SoP can be estimated on complete MSAs as well as pairwise projections (i.e. pairs of sequences extracted from the MSA they are part of while retaining the gaps). Here, the SoP was computed only on those residues that were assigned to the same structural state by DSSP^30^ while excluding loops.

### Transitive Consistency Score (TCS)

The TCS^15^ estimates the level of agreement between the alignment of any pair of sequences in an MSA and the realignment of this same pair across all possible sequence triplets within the same dataset. Triplets are estimated by combining all pairs of pairwise alignments featuring a common sequence. The method has been extensively described in ^15^ and has shown to be a robust predictor of sequence alignment accuracy. Given an MSA, the pairwise alignments required for the triplet analysis are generated by separately applying to every pair of sequences the same method used to generate the MSA being evaluated. This procedure is supported by the T-Coffee package in version v.13.45.57.844f401 and can be invoked by the following commands:

~~~
t_coffee -infile <MSA-Seq> -evaluate -method mafftginsi_pair
t_coffee -infile <MSA-AF2> -evaluate -method sap_pair,TMalign_pair
-pdb_dir <AF2>
~~~

### Kendall correlation analysis

Kendall correlation analysis quantifies the concordance between two different measures evaluated on the same samples. It estimates the frequency with which any increase of one measure is associated with an increase or decrease of the other measure. We applied this analysis on each of the structural accuracy measures (GDT-TS, pLDDT) and on the sequence accuracy predictor (TCS) in order to explore the agreement between these three measures, and the true sequence alignment accuracy as defined by the SoP. Since our datasets are independent from one another, we only included in the analysis the differences measured across pairs within families while ignoring differences across families. Kendall’s correlation (*τ*) coefficients were computed using :

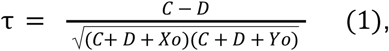

where *C* is the number of the concordant pairs within families, *D* is the number of the discordant pairs within families, *Xo* is the number of pairs tied only in the first variable and *Yo* is the number of pairs tied only in the second variable. Statistical significance was estimated by shuffling 1000 times the vector of the first variable, estimating the resulting *τ*, and comparing them to the true *τ* value. The *p*-value is then defined as the fraction of cases for which the true value was exceeded or equaled.

### Computation

All computation was carried out on a cluster running Scientific Linux release The structure predictions as well as the alignments, comparisons and evaluations were run within separate containers based on Ubuntu and Debian operating systems. The whole computational pipeline was implemented in the Nextflow language^31^ and was deployed in a containerized form using Singularity.

### Availability

All data, analyses, and results are available on Zenodo (https://doi.org/10.5281/zenodo.6564253). The code and scripts have been deposited in GitHub (https://github.com/cbcrg/msa-af2-nf) and the various containers in (https://cloud.sylabs.io/library/athbaltzis/af2/alphafold,https://hub.docker.com/r/athbaltzis/pred).

## Acknowledgements

We thank Guy Riddihough for revisions and comments on the manuscript.

## Author information

C.N., A.B. and L.M. designed the analysis, A.B. and L.M. carried out the validation and all the authors designed the validation procedure and wrote the manuscript.

