## supplements for "Improving sequence alignments with AlphaFold2 regardless of structural modeling accuracy"

This PDF file includes:

Supplementary Tables 1 to 4
Supplementary Figures 1 to 2

**Supplementary Table 1. Summary table with SoP scores per MSA algorithm per dataset using as reference the 3DCoffee alignments with the experimental structures (MSA-PDB).**

| Family | Number of sequences | Average length of sequences | FAMSA | G-INS-i (MSA-Seq) | MSAProbs | TCoffee | PSICoffee | MSA-AF2 | GM GDT-TS AF2 |
|---|---|---|---|---|---|---|---|---|---|
| **PF00001** | 16 | 359 | 71.70 | 74.10 | 80.40 | 83.00 | 85.30 | 97.70 | 75.09 |
| **PF00004** | 11 | 129 | 44.90 | 58.70 | 63.80 | 63.50 | 62.90 | 88.90 | 85.30 |
| **PF00073** | 17 | 169 | 81.50 | 87.20 | 89.60 | 88.70 | 82.80 | 100.00 | 93.82 |
| **PF00144** | 13 | 269 | 66.80 | 66.60 | 71.50 | 67.80 | 80.40 | 97.00 | 90.80 |
| **PF00271** | 12 | 112 | 72.50 | 89.50 | 89.40 | 89.50 | 91.10 | 96.40 | 92.68 |
| **PF00520** | 8 | 229 | 18.30 | 25.20 | 27.90 | 27.70 | 42.70 | 71.80 | 60.61 |
| **PF00664** | 13 | 266 | 62.30 | 68.20 | 54.80 | 61.70 | 88.40 | 98.20 | 65.30 |
| **PF00905** | 12 | 237 | 90.50 | 94.40 | 90.80 | 90.90 | 92.80 | 98.70 | 73.35 |
| **PF07992** | 10 | 259 | 64.10 | 72.80 | 71.80 | 68.80 | 76.40 | 97.10 | 96.08 |
| **PF12796** | 14 | 95 | 74.10 | 78.20 | 72.10 | 74.10 | 81.10 | 84.50 | 86.05 |
| **PF13306** | 11 | 100 | 56.10 | 49.60 | 59.90 | 73.20 | 60.90 | 99.60 | 93.26 |
| **PF13895** | 16 | 93 | 78.10 | 79.50 | 79.80 | 75.90 | 82.80 | 97.50 | 84.87 |
| **Average** | **12.75** | **193.08** | **65.08** | **70.33** | **70.98** | **72.07** | **77.30** | **93.95** | **83.10** |

**Supplementary Table 2. Comparison of the alignment accuracy of pairs of sequences between structure-based multiple sequence alignments using AlphaFold2 (MSA-AF2) models and sequence-based alignments (MSA-Seq) with a geometric mean of GDT-TS lower or greater than 75%.**

| | ***GDT-TS >= 75*** | ***GDT-TS < 75*** | ***Total*** |
|---|---|---|---|
| ***SoP(MSA-AF2) > SoP(MSA-Seq)*** | 563 (60.0%) | 172 (18.3%) | 735 |
| ***SoP(MSA-AF2) <= SoP(MSA-Seq)*** | 162 (17.3%) | 41 (4.4%) | 203 |

**Supplementary Table 3. Kendall analysis on SoP versus TCS, pLDDT and GDT-TS scores for each sequence within a family (Fig. 1c-e).**

| | **Concordant pairs** | **Discordant pairs** | **Tied pairs** | **Tau-b** | **p-value** |
|---|---|---|---|---|---|
| ***SoP vs TCS*** | 433 (46.2%) | 201 (21.4%) | 304 (32.4%) | 0.3 | p < 1e-03 |
| ***SoP vs pLDDT*** | 433 (46.2%) | 307 (32.7%) | 198 (21.1%) | 0.15 | p < 1e-03 |
| ***SoP vs GDT-TS*** | 425 (45.3%) | 314 (33.5%) | 199 (21.2%) | 0.13 | p < 1e-03 |

**Supplementary Table 4. NiRMSD scores per dataset per MSA algorithm computed using the experimental structures.**

| Family | FAMSA | G-INS-i | MSAProbs | TCoffee | PSIcoffee | 3D-Coffee_SAP +TMalign | 3DCoffee_TMalign | mTMalign |
|---|---|---|---|---|---|---|---|---|
| **PF00001** | 1.60 | 1.54 | 1.45 | 1.42 | 1.52 | 1.45 | 1.52 | 1.74 |
| **PF00004** | 3.71 | 3.02 | 3.08 | 2.93 | 2.86 | 2.02 | 1.98 | 1.92 |
| **PF00073** | 1.55 | 1.35 | 1.32 | 1.34 | 1.50 | 1.19 | 1.18 | 1.18 |
| **PF00144** | 1.19 | 1.21 | 1.26 | 1.26 | 1.12 | 0.97 | 0.96 | 0.89 |
| **PF00271** | 1.78 | 1.50 | 1.45 | 1.46 | 1.48 | 1.34 | 1.32 | 1.29 |
| **PF00520** | 6.04 | 5.50 | 5.30 | 5.65 | 3.55 | 2.84 | 3.01 | 3.18 |
| **PF00664** | 2.61 | 2.52 | 3.01 | 2.67 | 2.11 | 1.96 | 1.99 | 2.40 |
| **PF00905** | 1.25 | 1.20 | 1.19 | 1.21 | 1.19 | 1.07 | 1.06 | 0.98 |
| **PF07992** | 1.76 | 1.62 | 1.58 | 1.73 | 1.59 | 1.22 | 1.24 | 1.20 |
| **PF12796** | 1.49 | 1.52 | 1.53 | 1.53 | 1.41 | 1.18 | 1.21 | 1.18 |
| **PF13306** | 1.18 | 1.29 | 1.25 | 1.45 | 1.23 | 1.03 | 1.15 | 1.03 |
| **PF13895** | 1.91 | 1.77 | 1.72 | 1.79 | 1.66 | 1.30 | 1.33 | 1.29 |
| **Average** | **2.17** | **2.00** | **2.01** | **2.04** | **1.77** | **1.46** | **1.50** | **1.52** |

a

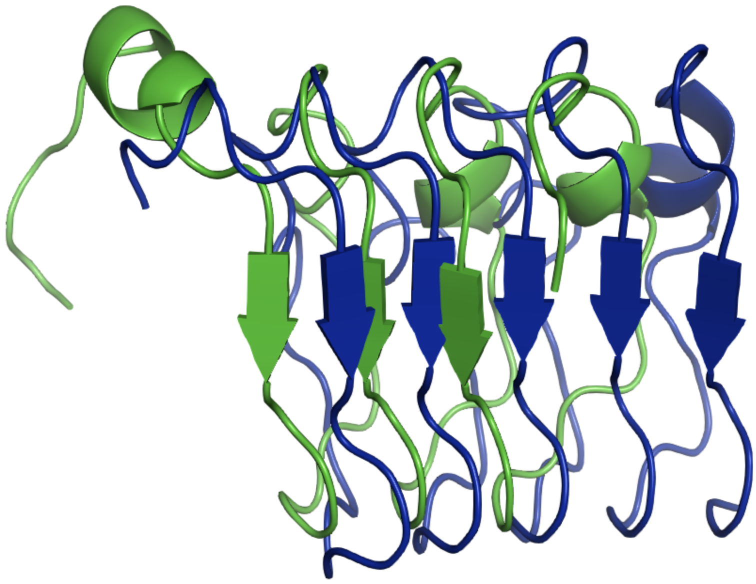

b

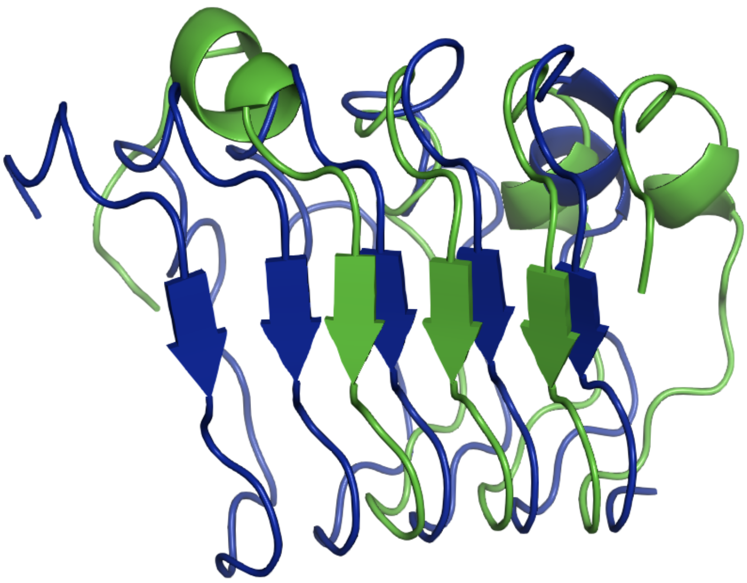

**Supplementary Fig. 1 | Impact of repeated elements on sequence-based and structure-based alignments. a,** Sequence-based alignment of the PDB entries 6u12a_30-111 and 6tl8a_53-165 that belong to the PF13306 dataset performed by PyMOL “align” algorithm. **b,** Structure-based alignment of the PDB entries 6u12a_30-111 and 6tl8a_53-165 that belong to the PF13306 dataset performed by PyMOL “super” algorithm.

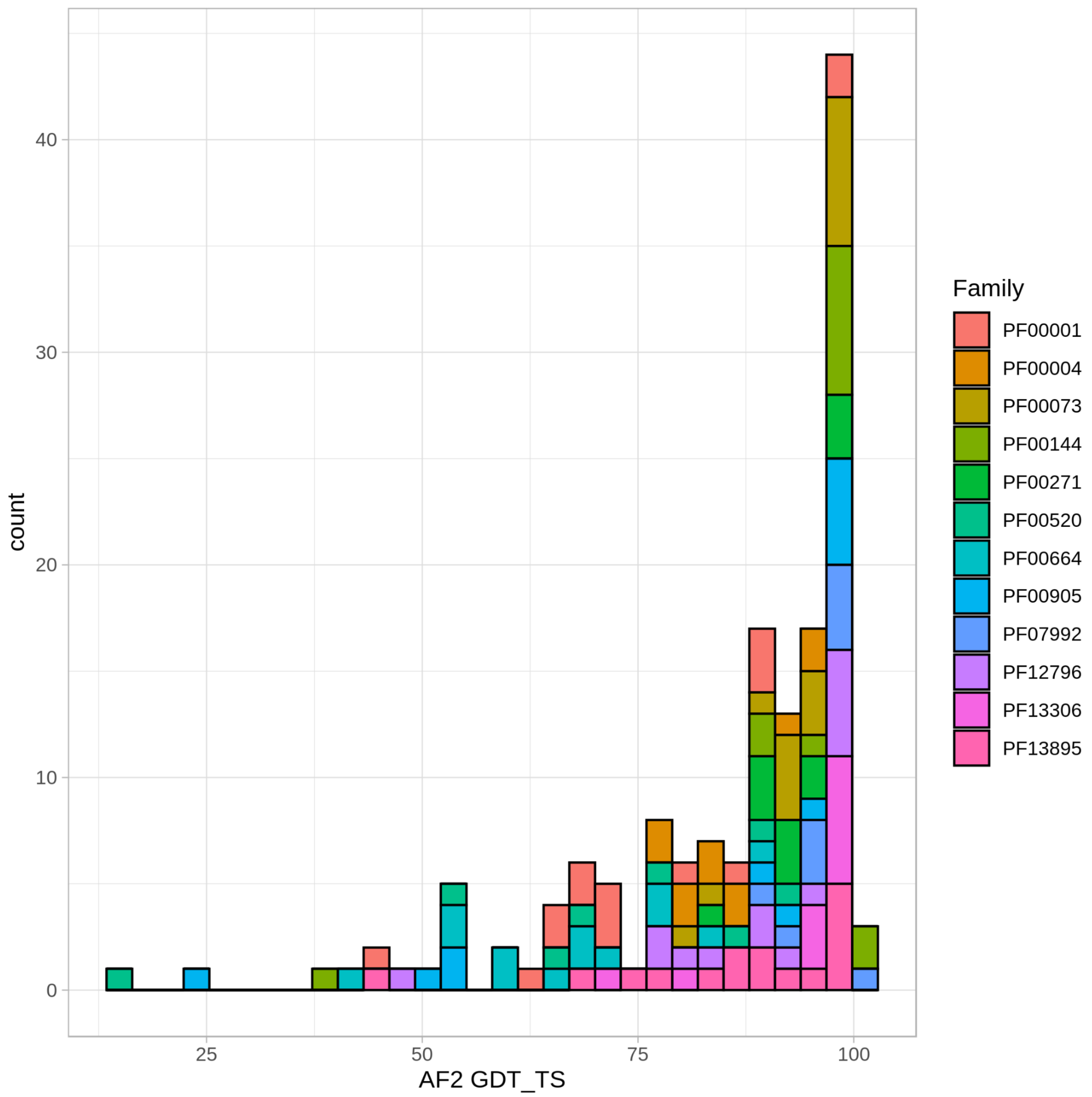


**Supplementary Fig. 2 | Distribution of structural correctness of the AlphaFold2 models.** Histogram of GDT-TS scores between the AF2 predicted and the equivalent experimental structures colored by dataset.
